# Orally consumed cannabinoids provide long-lasting relief of allodynia in a mouse model of chronic neuropathic pain

**DOI:** 10.1101/556373

**Authors:** Edward J.Y. Leung, Antony D. Abraham, Brenden A. Wong, Lauren C. Kruse, Jeremy J. Clark, Benjamin B. Land

## Abstract

Chronic pain affects a significant percentage of the United States population, and available pain medications like opioids have drawbacks that make long-term use untenable. Cannabinoids show promise in the management of pain, but long-term treatment of pain with cannabinoids has been challenging to implement in preclinical models. We developed a voluntary, gelatin oral self-administration paradigm that allowed animals to consume Δ9-tetrahydrocannabinol, cannabidiol, or morphine *ad libitum*. Animals stably consumed these gelatins over 3 weeks, with detectable serum levels. We designed a real-time gelatin measurement system, and observed that mice consumed gelatin throughout the light and dark cycles, with THC-gelatin animals consuming less than the other groups. Consumption of all three gelatins reduced measures of allodynia in a chronic, neuropathic sciatic nerve injury model, but tolerance to morphine developed after one week while THC or CBD reduced allodynia over three weeks. Hyperalgesia took longer to develop after sciatic nerve injury, but by the last day of testing THC significantly reduced hyperalgesia responses, with a trend effect of CBD, and no effect of morphine. Mouse vocalizations were recorded throughout the experiment, and mice showed a large increase in ultrasonic, broadband clicks after sciatic nerve injury, which was reversed by both THC and CBD. This study demonstrates that mice will voluntarily consume both cannabinoids and opioids via gelatin, and that cannabinoids can provide long-term relief of chronic pain states. Additionally, ultrasonic clicks may objectively represent the pain status of a mouse and could be integrated into future pain models.

## Introduction

Chronic pain, defined as pain that lasts at least 12 weeks, affects between 40-100 million individuals in the United States (Johannes et al, 2010; Committee on Advancing Pain Research, 2011; Pitcher et al, 2019). The severity of chronic pain can differ, but approximately 11 million adults report having major restrictions in activity due to their pain state (Pitcher et al, 2019), and the total national economic burden per year is estimated to be approximately $600 billion (Gaskin and Richard, 2017). Current treatments for chronic pain are largely limited to non-steroidal anti-inflammatory drugs (NSAIDS) and opioids (opiates), each with significant drawbacks. NSAIDS can produce toxic side effects in the kidney (Lee et al, 2007), and thus high or frequent doses are unachievable. Opioids, including morphine and its derivatives like oxycodone, produce analgesic tolerance rapidly (Williams et al, 2013; Schattauer et al, 2017), requiring escalation of dose to achieve pain relief over time. Furthermore, opioids produce euphoria in patients and are rewarding in rodent models (Fields and Margolis, 2015). The combined effects of tolerance and reward can lead to opioid dependence and substance use disorder. The large increase in prescriptions for opioid medications indicated for pain over the last 20-30 years has contributed to the current opioid crisis in the United States (Phillips et al, 2017), and thus it is important to identify and validate alternative medications for chronic pain treatment.

Cannabinoids are promising candidates for the treatment of pain (Manzanares et al, 2006; Elikottil et al, 2009; Vučković et al, 2018). Numerous studies in preclinical models and human trials have shown that the principal cannabis phytocannabinoids Δ9 tetrahydrocannabinol (THC) and cannabidiol (CBD), as well as synthetic cannabimimetic ligands, are able to relieve various pain states, although the extent to which they provide true pain relief in humans remains controversial (Romero-Sandoval et al, 2018; Lötsch et al, 2018). These pain states include measures of acute analgesia, as well as neuropathic, inflammatory, and cancer pain modalities (Elikottil et al, 2009; Xiong et al 2011; Rock et al, 2019). Likewise, a large number of receptor systems have been implicated in mediating cannabinoid analgesia, including the cannabinoid receptors themselves (Mulpuri et al, 2018; Li et al 2019; Grenald et al, 2017), TRP channels (Muller et al, 2019), G protein-coupled receptors (Rodríguez-Muñoz et al, 2018; Guerrero-Alba et al, 2019), and glycine receptors (Xiong et al, 2011, 2012). While many of these studies have focused on acute outcomes of only one or several drug administrations, long-term studies of cannabinoids in humans have shown mixed, and often subtle effects in treatment of pain (Vučković et al, 2018; Lötsch et al, 2018), suggesting that acute pain models may be inadequate when considering chronic pain. To better predict treatment outcomes in humans, we aimed to develop a preclinical model for long-term cannabinoid intake in the context of chronic pain states.

One major challenge to long-term administration of cannabinoids in preclinical models is that most have historically relied on experimenter administration, which does not closely resemble intake patterns of humans in either route of administration, dose, or timing. Efforts have recently been made to model human cannabinoid consumption in rodents more closely, including use of vapor chambers (Nguyen et al, 2016; Javadi-Paydar et al 2018), or sweetened, orally consumed solutions (Barrus et al, 2018). However, in these models, intake was still limited to a relatively short period or was not voluntary. To gain a better understanding of cannabinoid consumption for long-term pain management, we adapted an *ad libitum* gelatin, oral self-administration paradigm (Schindler et al, 2014; Nasrallah et al, 2011) for the cannabinoids THC and CBD. Using this model in mice, we examined long-term (3 week) consumption patterns and feeding microstructure. We then assessed the long-term efficacy of these two compounds in reducing chronic, neuropathic pain following partial sciatic nerve ligation in comparison with orally consumed morphine.

## Methods

### Animals

Male and female C57BL/6 mice ranging from 2 to 4 months of age were used in these experiments. All animals were singly housed under normal conditions (dark cycle beginning at 8:00 pm and ending at 6:00 am), with standard chow and water *ad libitum*. All testing was performed from 1-4 pm. All animals were drug naive with no prior procedures performed. Animal procedures were approved by the Institutional Animal Care and Use Committee of the University of Washington and conform to the guidelines on the care and use of animals by the National Institutes of Health.

### Drugs

Δ9-tetrahydrocannibinol (THC), cannabidiol (CBD), and morphine were provided by the National Institute of Drug Abuse Drug Supply Program (Bethesda, MD). THC in ethanol (EtOH) (100 mg/ml) was diluted using 95% EtOH to a stock concentration of 20 mg/ml. CBD was dissolved into 95% EtOH to a stock concentration of 20 mg/ml. For i.p. injections, cannabinoids were first dissolved in 5% EtOH, then 5% cremophor was added, followed by sterile saline. Morphine was dissolved in sterile saline.

### Gelatin

3.85 g of Knox Gelatin was dissolved into 100 ml of deionized H_2_O and raised to a temperature of 40^°^C. Then either 5 g, 2.5 g or 1 g of Polycal sugar was dissolved into the mixture to make the 5%, 2.5% and 1% sugar content gels respectively. While maintaining the temperature of the mixture between 41-43^°^C, THC or CBD (20 mg/ml) was added to achieve a final drug concentration of 1 mg/15 ml (g) of gelatin. Morphine was added to achieve a concentration of 1.125 mg/15 ml (g). Gelatin was allowed to cool in 4^°^C refrigerator before use.

### Oral gelatin self-administration

Animals were singly-housed, and a small plastic cup was secured to the metal wire top using a hose clamp. Approximately 10 g of control gelatin (no drug) at 5% Polycal content was inserted into each of the plastic cups at approximately 2:00 pm for 3 days. The weight of the gelatin from the previous day was recorded and the remaining gelatin was discarded. After this, the Polycal content was reduced to 2.5% for two days, and again to 1% for 5 days. Two days prior to sciatic nerve ligation, gelatins were removed, and gelatin with drug was introduced one day after ligation, and every day thereafter until the study was concluded.

### Measurement of feeding microstructure

A piezoelectric load cell was attached to a 3-D printed frame including the gelatin cup. The signal from the load cell was amplified and then sent to an Arduino mounted above the top of the cage for processing using custom code, and data was then stored on a flash storage device. Disturbances were defined as changes in mass greater than 0.04 g between recorded samples (2 s sample rate). A disturbance, or series of disturbances, was classified as a bout if there was a period greater than 120 seconds between consecutive disturbances. Schematics and all source code is available at GitHub: https://github.com/bwong07/continuous-gel-self-administration-device

### Partial sciatic nerve ligation (pSNL)

pSNL was performed as previously described (Seltzer et al, 1990; Xu et al, 2004). Briefly, mice were anesthetized using isoflurane, and the sciatic nerve was exposed approximately halfway between the hip and knee using a scalpel. A silk suture was then passed through approximately 1/2 of the nerve bundle, before being tightly ligated and crushed. The fascia and skin were then sutured closed.

### Hotplate

A hotplate apparatus (IITC Life Science) was set to a temperature of 57.5^°^C. Mice were gently placed onto the hotplate, and the latency of a pain response was recorded. A pain response was defined as either a paw lick, jump or a hind paw shuffle.

### Von Frey

Von Frey hairs (IITC Life Science) of various, predetermined weights were used (0.1g – 8g). Individual hairs of increasing weight (force) were pressed against each hind paw until a response (defined as a hind paw lick, hind paw lift or a hind paw jump) was observed. The lowest weight that elicited a response for 2 out of 3 presses was recorded. We observed no differences in the ipsilateral and contralateral paw withdrawal scores (Koltzenburg et al, 1999; Xu et al, 2004; Arguis et al, 2008), so the two paws were averaged for each mouse.

### pSNL pain testing schedule

Mice were tested for hyperalgesia and allodynia (pain test) on the hotplate and von Frey, respectively. Baseline responses were measured 3 days prior, and immediately before surgery. A third pain test was then administered 24 hours after pSNL (day 1 post-pSNL), confirming a neuropathic pain state. Pain tests were then administered immediately prior to gelatin insertion at approximately 2:00 pm on days 5-, 7-, 12-, 15-, 18- and 22-post pSNL. Pain data is expressed as percentage of baseline, which was calculated by averaging the two pre-pSNL pain test scores and dividing each subsequent pain test post-pSNL by that baseline value. Percentages below 100 indicate increases in pain.

### Acute pain testing schedule

Mice were given control, THC, CBD, or morphine gelatins, as described above, for 7 days. On day 7, an initial pain test was administered for all groups. On day 8, initial rectal body temperatures were taken for control, THC, and CBD groups, and mice were then injected with vehicle, THC (30 mg/kg), CBD (30 mg/kg), or morphine (10 mg/kg), corresponding to their drug gelatin. 30 min after injection, pain tests were administered for morphine-treated mice. 2 h after control, THC, or CBD injection, rectal body temperatures were recorded, and pain tests were and administered. Drug-naïve animals went through the same procedure, with no gelatin exposure.

### Ultrasonic Vocalizations

A Pettersson microphone (Norway, model M500-384, maximum recording frequency = 192 KHz) was inserted through a small hole in a filter top placed on top of the mouse’s home cage. 5-minute recordings were taken immediately before and after a pain test session using Avisoft SASLab Lite recording software. Recordings were analyzed using RavenLite (Cornell lab of Ornithology), by first generating a spectrogram of the recording and then manually scanning each file for ultrasonic broadband clicks, defined as an audio feature that met several criteria: 1) Lasting less than 10 ms, 2) Having a minimum 10 KHz frequency width, 3) Having an average power density of greater than −50 dB FS over 5 ms at its minimum frequency width, 4) Having peak power in the ultrasonic range (20 KHz to 100 KHz), 5) Lacking a fundamental frequency or appreciable power below 15 KHz. These last two criteria were used to reject audible clicks. The number of clicks in each 5 min recording were quantified.

### Serum levels of drugs

Animals were rapidly decapitated and trunk blood was collected, incubated at RT for 15-20 min, and spun for 10 min at 1800 x g, at 4^°^C for serum. Serum was stored at −80^°^C and sent to the Mass Spectrometry Facility at University of Washington for processing against known standards for THC, CBD, and morphine.

### Data Analysis

All data were analyzed using MATLAB and GraphPad Prism 7-8. For all statistical testing (paired t-test, One- and Two-way ANOVA, and relevant post-hoc testing), alpha level was set to 0.05.

## Results

Based on the gelatin free-access model developed by Clark et al (Schindler et al, 2014; Nasrallah et al, 2011), we designed a paradigm that would allow us to study long-term treatment outcomes of cannabinoid and opioid self-administration in mice with chronic neuropathic pain (**Figure 1a**). After both sucrose fading and drug introduction, gelatin consumption was consistent within groups across the experiment (**Fig. 1b**), with no significant differences in consumption across days of access in any group (2-way ANOVA, control gelatins main effect of time F_4,180_ = 1.743, p = 0.14; drug gelatins main effect of time F_16,720_ = 1.342, p = 0.16). Consumption of gelatin containing THC was significantly lower compared to control after pSNL (main effect of treatment F_3,45_ = 231, p = 0.003, Dunnett’s Control vs THC p = 0.012), but this effect was not observed in either the CBD or morphine gelatins (Control vs CBD p = 0.45, Control vs Morphine p = 0.96). Similarly, there was no difference in total gelatin consumed throughout the 3 weeks after pSNL in CBD and morphine groups (**Fig. 1c,** 1-way ANOVA F_3,43_ = 5.92, p = 0.0018, Dunnett’s Control vs CBD p = 0.71, Control vs Morphine p = 0.84), but THC gelatin consumption was significantly lower than control (Control vs THC p = 0.005). There was no apparent escalation of consumption following initial introduction of drug for any of the gelatins. To understand if the decrease in THC gelatin consumption was due to the presence of THC or the neuropathic pain state, a separate cohort of animals were trained to consume either THC or CBD gelatin both before and after pSNL (**Supplemental Figure 1**). In both THC and CBD groups, the level of gelatin consumption after pSNL remained the same, indicating that occurrence of chronic pain does not alter consumption or produce escalated intake (**Fig. 1d**, 1-way ANOVA THC F_1.45,8.72_= 1.79, p = 0.22; CBD F_1.65,11.55_= 0.99, p = 0.39). To confirm that consumption of the drug gelatin produced circulating levels of drug in the mice, trunk blood was collected either 7 or 24 hours after the last gelatin insertion and serum was prepared. Detectable levels of all drugs were observed at every time point measured (**Fig. 1e**). Drug levels in THC- and CBD-consuming mice were similar between the 7 and 24 h time points, suggesting that animals maintain a relatively stable blood level of drug throughout the day. This prompted us to explore the patterns of drug self-administration throughout the period of drug availability.

**Figure 1.**
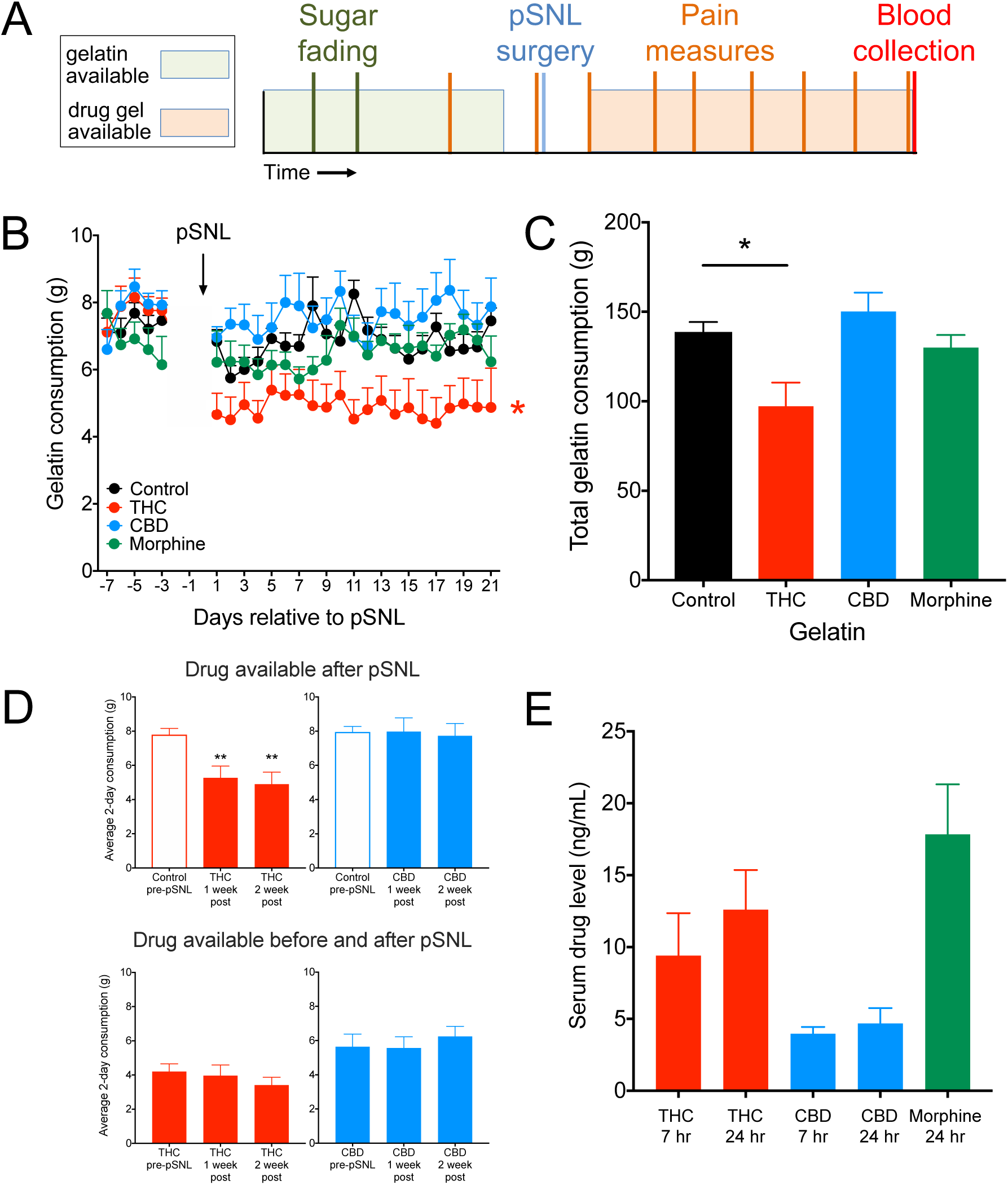
Mice stably consume cannabinoid and opioid gelatin over 3 weeks. **A**) Schematic of study design for chronic neuropathic pain and gelatin consumption. **B**) Average daily consumption of gelatin for the 4 groups (control, THC, CBD, and morphine, n = 10-17). Prior to pSNL, all groups consumed control gelatin. Mice consuming THC-gelatin ate significantly less than control mice (p = 0.003). **C**) Total consumption of gelatin post-pSNL for control, THC, CBD, and morphine gelatins. Total THC-gelatin consumption was significantly lower than control mice (p = 0.005). **D**) Comparisons within groups between the average consumption over the last two days prior to pSNL, days 6-7 after pSNL, and days 13-14 after pSNL. The top row represent animals who only received THC or CBD after pSNL. The bottom row represent animals that received THC or CBD before and after pSNL. Animals receiving drug gelatin prior to pSNL showed no differences in consumption after pSNL. **E**) Serum levels of THC, CBD, and morphine collected either 7 or 24 h after the final gelatin insertion (n = 2-6). * < 0.05, ** < 0.01.

To enable in-depth analysis of drug consumption, we designed a novel measurement device that was able to track the weight of the gelatin and any disturbances of the gelatin cup in real-time. The device consisted of a piezoelectric sensor (load cell) that measured changes in mass, a signal amplifier, and a microcontroller connected to a data storage device (**Figure 2a, Supplemental Figure 2a**). Mouse interactions with the gelatin cup, which registered as brief and rapid changes in mass (“disturbances”), were recorded, as well as the overall, longer term changes in gelatin mass. A representative trace of both gelatin cup disturbances and cumulative gelatin consumed showed frequent interactions with gelatin cup and concordant increases in gelatin intake (**Fig. 2b**). This relationship was confirmed when we correlated the number of feeding bouts (defined as one or more disturbances separated from preceding or following disturbances by greater than 120 s) with the amount of gelatin consumed in a 2 h period (**Fig. 2c**, linear regression r^2^ = 0.44, F = 98.5, p < 0.0001). Using this device to record the consumption patterns for each type of gelatin (**Fig. 2d**), we observed stable consumption of all gelatins throughout the recording period, and THC consumption was significantly lower than control by the end, as in Figure 1 (1-way ANOVA, F_3,10_ = 5.70, p = 0.015, Dunnett’s Control vs THC p = 0.044). However, there were no significant differences between any of the groups in the number of feeding bouts throughout the day (**Fig. 2e**). This suggests that mice receiving THC gels were not feeding less frequently, but were consuming less during each bout. Gelatin consumption followed expected circadian patterns, and consumption was higher during the dark phase when the animals were awake and more active (**Supplemental Figure 2b**, 2-way ANOVA, main effect of time, F_1,10_ = 43.04, p < 0.001, Holm-Sidak, all groups, Dark cycle vs Light cycle p < 0.05).

**Figure 2.**
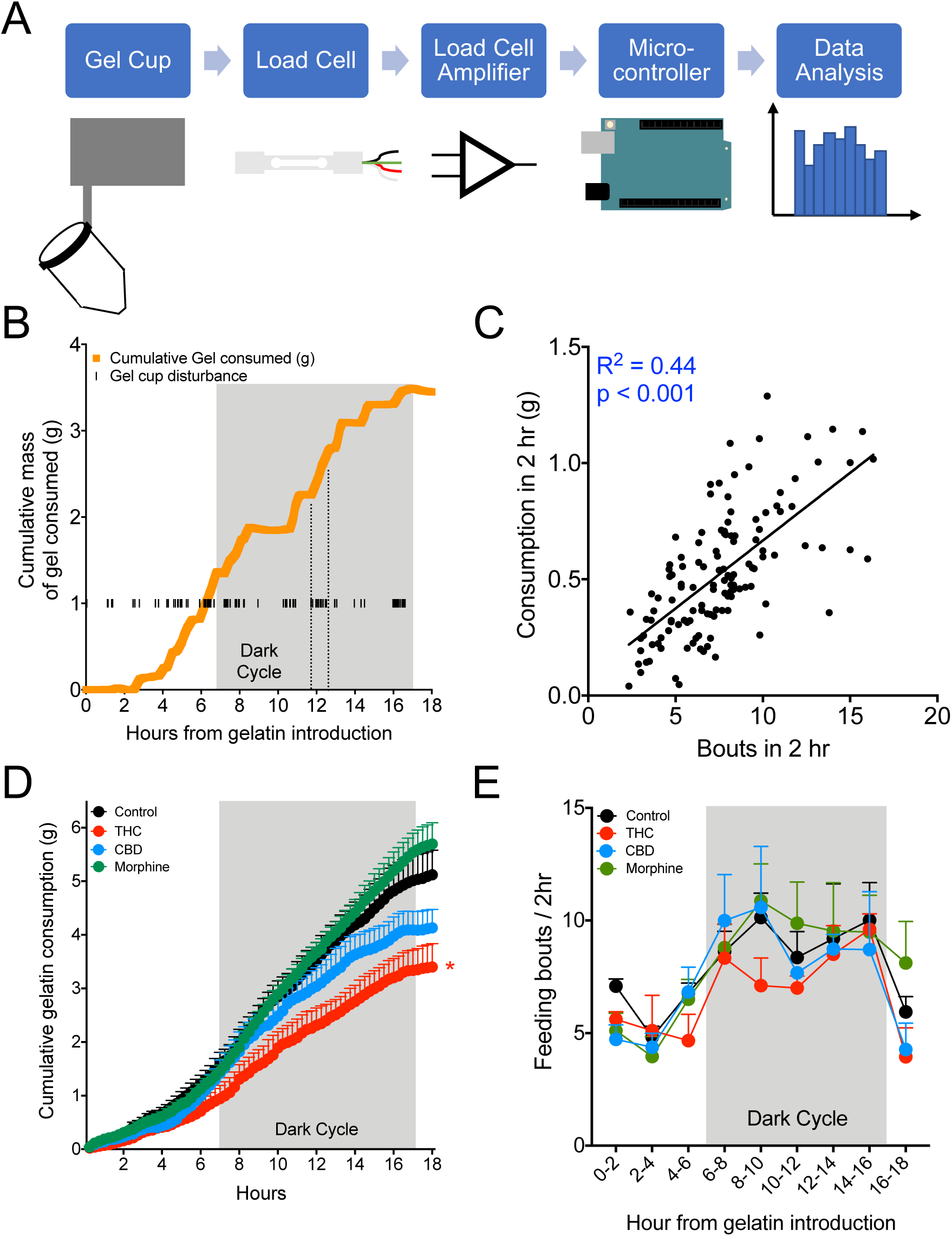
Microstructure of gelatin consumption. **A**) Schematic and signal flow of piezoelectric load cell that measures second-by-second consumption and disturbances of the gelatin cup. **B**) Example of cumulative gelatin consumption and disturbances of the gelatin cup over 18 h from one mouse. **C**) Correlation between the number of disturbances of the gelatin cup and the consumption of gelatin in 5 min windows (n = 126 points from 14 animals, p < 0.001). **D**) Average cumulative consumption of the control, THC, CBD, and morphine gelatins over the 18 h measurement period. By 18 h, THC mice showed significantly less cumulative consumption than controls (n = 3-4 mice per group, p = 0.04). **E**) Average number of bouts, defined as a set of disturbances preceded or succeeded by at least two minutes, during 2 hour windows throughout the measurement period. There are no differences between the groups in the number of bouts (n = 3-4 mice per group). * p < 0.05.

Once we confirmed that mice were stably consuming gelatin, we tested to see if this level of consumption was sufficient to produce significant relief of chronic neuropathic pain. The pSNL surgery produced allodynia, a painful response to a normally non-painful stimulus as measured by Von Frey hairs, in mice after one day (**Figure 3a**). This allodynia persisted throughout the experiment in animals that only received control gelatin, and both THC and CBD significantly relieved allodynia compared to control mice (**Fig. 3a**, 2-way ANOVA, interaction F_15,235_ = 3.07, p < 0.001, Holm-Sidak post-hoc). This effect was apparent on the first pain test after drug-gelatin presentation (day 5) and was maintained throughout the 3 weeks of testing, with a modest decrease in THC efficacy on the final two testing days. Morphine relieved allodynia maximally on the sixth day following initial introduction of drug (day 7), but on day 12 post-pSNL paw withdrawal response thresholds decreased to similar levels as control animals. Examining only the final day of testing, mice that received either THC or CBD gelatin had significantly reduced allodynia compared to mice that received control gelatin (**Fig. 3b**, 1-way ANOVA, F_3,47_ = 5.885, p = 0.002, Holm-Sidak Control vs. THC p = 0.049, Control vs. CBD p = 0.002). There was no difference between mice that received morphine or control gelatin (Control vs Morphine p = 0.96). In contrast to the clear effects of cannabinoids on allodynia, the hyperalgesia (an increased pain response to a normally painful stimuli) measured by hot plate took longer to develop following pSNL and was more variable in all groups (**Fig. 3c**). In the first week of drug gelatin exposure, morphine produced a significant analgesic effect that disappeared by day 12 post-pSNL, while neither THC or CBD produced any significant effect (2-way ANOVA, interaction F_15,160_ = 1.76, p = 0.044, Holm-Sidak post-hoc). Examining only the final day of testing (**Fig. 3d**), THC showed a significant reduction in hyperalgesia compared to controls, while there was a trend effect of CBD and no effect of morphine (1-way ANOVA, F_3,32_= 3.2, p = 0.036, Holm-Sidak Control vs THC p = 0.02, Control vs CBD p = 0.093, Control vs Morphine p = 0.35).

**Figure 3.**
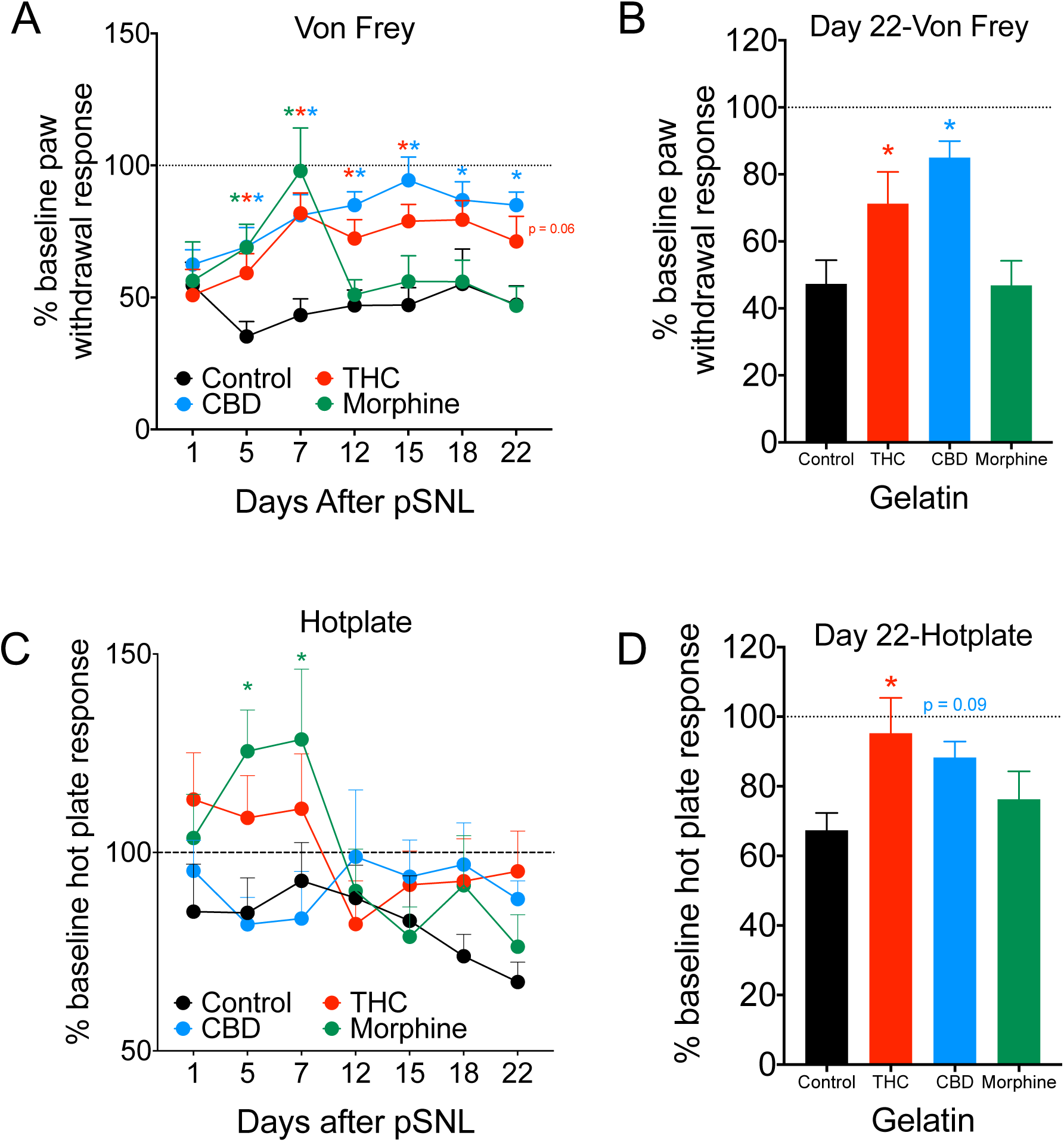
Cannabinoid consumption decreases measurements of neuropathic pain over 3 weeks. **A**) Paw withdrawal thresholds, plotted as a percentage of pre-pSNL response, are shown 1 day after pSNL, and on the 7 subsequent testing days after introduction of either control, THC, CBD, or morphine gelatin. THC and CBD groups showed long-lasting reduction in allodynia, while the effect of morphine lasted approximately 1 week (n = 10-19 per group). **B**) Comparison of percentage of pre-pSNL response on day 22, the final day of pain testing, between the 4 treatment groups (n = 10-19 per group). **C**) Latency to respond to hotplate, plotted as a percentage of pre-pSNL response, are shown 1 day after pSNL, and on the 7 subsequent testing days after introduction of either control, THC, CBD, or morphine gelatin. The morphine group showed an analgesic response in the first week, while no other groups were significantly different from control (n = 8-12 per group). **D**) Comparison of percentage of pre-pSNL response on day 22. THC significantly reduced hyperalgesia while there was a trend effect of CBD (n = 8-12 per group). * p < 0.05.

To look at analgesic responses in a neuropathic pain-naïve state and to confirm the development of morphine tolerance, we next examined behavioral effects of experimenter-administered drug in drug-naive and drug-exposed animals with 7 days of gelatin self-administration. Consumption of either THC or CBD gelatin for 7 days did not significantly change basal measurements of body temperature, analgesia (hotplate response), or allodynia (Von Frey) from mice receiving control gelatin (**Figure 4a,c,e**). Mice that consumed morphine gelatin also showed no significant differences from control gelatin for analgesia or allodynia (**Fig. 4c,e**). We then tested whether prior gelatin consumption produced tolerance by injecting the same mice with their corresponding drugs and comparing their behavioral scores with drug-naïve mice receiving a matching injection. As expected, THC injection (30 mg/kg i.p.) produced a significant decrease in body temperature, increased paw withdrawal threshold, and increased hot plate analgesia compared to controls (**Fig. 4b,d,f**, 2-way ANOVAs, body temperature interaction F_2,18_ = 4.72, p = 0.023, Control vs THC < 0.001; Von Frey F_3,23_= 6.63, p = 0.002, Control vs THC = 0.006; hotplate F_3,28_= 3.20, p = 0.038; Holm-Sidak treatment differences Control vs THC p < 0.001). In contrast, CBD injection did not change any of the behavioral measurements compared to control or drug-naïve mice. This was surprising and suggests that CBD may only produce an analgesic effect if pain is already established. There were no significant differences between 7-day gelatin and drug-naïve animals on any measure for THC or CBD (**Fig. 4 b,d,f**). Finally, morphine injection in drug-naïve mice produced significant increases in analgesia and paw withdrawal threshold (**Fig. 4 d,f**, Holm-Sidak drug-naïve Von Frey Control vs Morphine p < 0.001, hotplate Control vs Morphine p = 0.007,). However, these effects were not present in animals that had consumed morphine gelatin for 7 days, verifying our previous observation that mice had developed tolerance to morphine (Holm-Sidak Morphine naïve vs 7-day, Von Frey p < 0.001, hotplate p = 0.037,). These results demonstrate that week-long consumption of THC, in contrast to morphine, does not produce tolerance.

**Figure 4.**
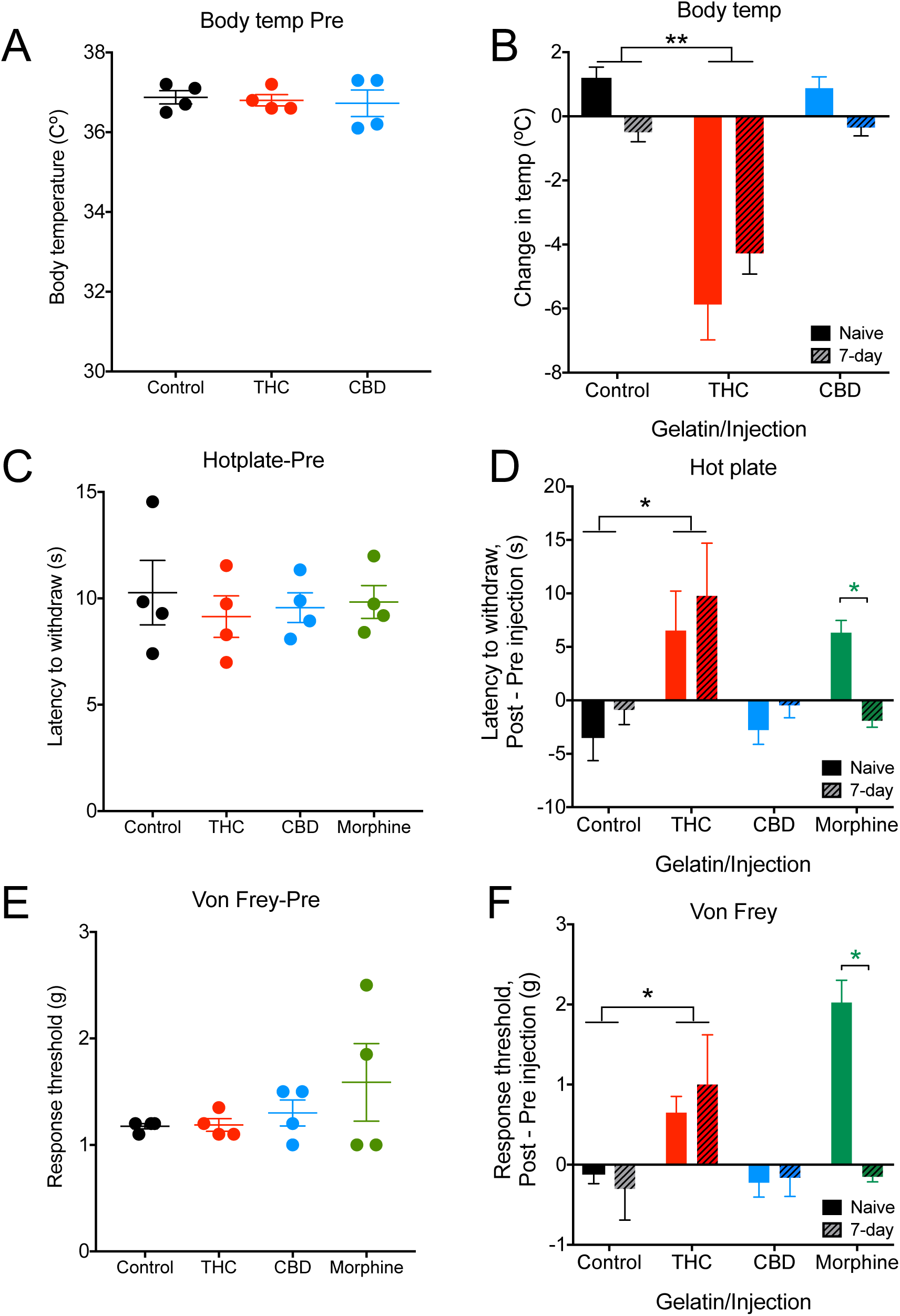
Measures of analgesia and body temperature in neuropathic pain-naïve mice. **A**) Body temperature measured after 7 days of control, THC, or CBD gelatin consumption (n = 4 per group). **B**) Change in body temperature 2hr after either vehicle, THC (30 mg/kg i.p.), or CBD (30 mg/kg i.p.) experimenter-administered dose in 7-day gelatin consuming animals or drug-naïve controls. THC injected mice showed the predicted decrease in body temperature in both groups, whereas CBD showed no change (n = 4 per group). **C**) Baseline pain responses in the hotplate after 7 days of control, THC, CBD, or morphine gelatin consumption (n = 4 per group). **D**) Change in latency to pain response 2hr after either vehicle, THC (30 mg/kg i.p.), CBD (30 mg/kg i.p.), or 30 min after morphine (10 mg/kg i.p.) experimenter-administered dose in 7-day gelatin consuming animals or drug-naïve controls. THC injected mice in both groups showed a similar increase in analgesic response, while CBD animals showed no change. Drug-naïve morphine injected animals showed analgesia, while there was no change in mice that have consumed morphine for 7 days (n = 4-8 per group). **E**) Baseline allodynia response using Von Frey hair after 7 days of control, THC, CBD, or morphine gelatin consumption (n = 4 per group). **F**) Change in paw withdrawal threshold 2hr after either vehicle, THC (30 mg/kg i.p.), CBD (30 mg/kg i.p.), or 30 min after morphine (10 mg/kg i.p.) experimenter-administered dose in 7-day gelatin consuming animals or drug-naïve controls. THC injected mice in both groups showed a similar increase in paw withdrawal threshold, while CBD animals again showed no change. Drug-naïve morphine injected animals showed an increase in paw withdrawal threshold, while there was no change in mice that have consumed morphine for 7 days (n = 4 per group). * p < 0.05, ** p < 0.01.

Because measurements of pain in mice often require a subjective scoring metric, we explored whether mice exhibited different types of vocalizations as a function of chronic neuropathic pain and subsequent treatment. This could provide a less subjective readout of the animals’ pain status. Using a wide frequency range microphone (up to 192 KHz, Petterson), we recorded ultrasonic vocalizations of mice in their home cages 5 min before pain testing on 4 days in control, THC, and CBD gelatin groups. In these singly-housed mice, we did not observe many complex vocalizations, such as rising or falling calls. However, we did notice the presence of many ultrasonic, broadband clicks, a mouse vocalization that has been previously hypothesized to be a negative con-specific signal (Sugimoto et al, 2011; **Figure 5a, Supplemental Figure 3a**). In a neuropathic pain-free state, mice showed a significant increase in ultrasonic clicks after pain tests were administered (**Supplemental Fig. 3b**, paired t-test t_5_ = 2.64, p = 0.046). These clicks were not attributable to other environmental stimuli, as empty cage recordings show very few clicks (**Supplemental Fig. 3b**). Compared to their neuropathic pain-free state, mice significantly increased click vocalizations following pSNL surgery (**Fig. 5b**, paired t-test t_15_ = 6.14, p < 0.001). Furthermore, 4-days after initial introduction of either THC or CBD gelatin, click vocalizations were significantly reduced compared to control gelatin, and were comparable to baseline counts prior to pSNL surgery (**Fig. 5b**, 1-way ANOVA F_2,13_= 14.46, p < 0.001, Dunnett’s Control vs THC p < 0.01, Control vs CBD p < 0.01). Because of variability in click vocalizations prior to the introduction of drug, the same data was separated by treatment group (**Fig. 5c**). Although there was variability in the magnitude of response (driven largely by the large variance in clicks after neuropathic pain), the effect of THC or CBD in reducing the number of clicks remains significant (**Fig. 5c**, 1-way ANOVA, all groups < 0.05, Tukey’s post-hoc < 0.05).

**Figure 5.**
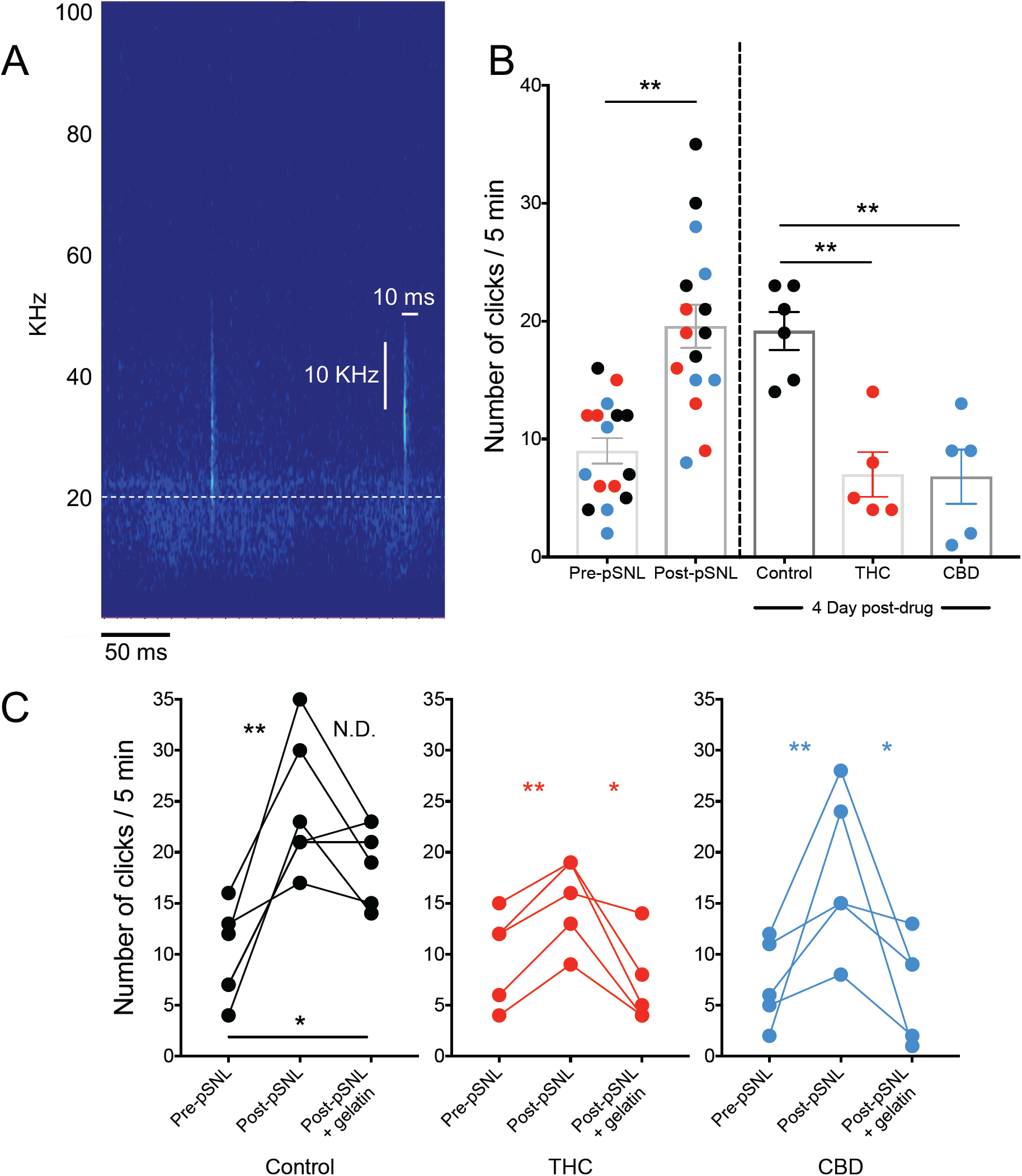
Ultrasonic click vocalizations track pain state. **A**) Two representative ultrasonic clicks. Each is under 10 ms, wider than 10 KHz, with no appreciable power in the audible range. Dashed line represents 20 KHz, the upper limit of the human audible range. **B**) Number of ultrasonic clicks per 5 min recording interval. Colors represent control (black), THC (red), and CBD (blue) gelatin mice (n = 5-6 per group). pSNL significantly increased the number of clicks (p < 0.001). After 4 days of gelatin, the number of clicks remained elevated in control mice, while THC and CBD mice showed decreases similar to baseline counts. **C**) Separated clicks for each treatment group. All groups showed a significant increase in clicks after pSNL, and in THC and CBD consuming mice clicks significantly decreased after drug-gelatin consumption. * p < 0.05, ** p < 0.01.

## Discussion

The major findings of this study are that 1) Animals will voluntarily consume both cannabinoids and opioids in a gelatin preparation, 2) These doses are sufficient to produce analgesia over short (morphine) and long (THC and CBD) time periods without dose escalation, and 3) Ultrasonic clicks may be an ethological and objective representation of pain status. Importantly, these findings indicate that both THC and CBD may be suitable for long-term treatment of chronic pain, in contrast to morphine. CBD may be especially beneficial in treatment of chronic pain because it lacks the rewarding psychoactive effects of THC and does not produce analgesia in a pain-naïve state (Figure 4), while still providing significant relief of allodynia during chronic pain. Hyperalgesia and allodynia reflect different environmental stimuli, and while predictable painful stimuli can often be avoided, allodynic responses to stimuli like normal touch cannot. Therefore, the long-term inhibition of allodynia by CBD, without tolerance or psychoactive side-effects, could be a future indication along with its established role in seizure prevention (Kaplan et al, 2017). Why CBD fails to produce analgesia in pain-naïve mice is unknown, but may be related to its established anti-inflammatory role (Li et al, 2018, Mukhopadhyay et al, 2011), and that pro-inflammatory pain signals are required for CBD to be effective.

The use of an oral gelatin-based model of self-administration offers several benefits over existing methods for cannabinoid and opioid intake. Animals are able to titrate their dose by increasing or decreasing consumption, and there is a negligible caloric component. We saw evidence of this titration in the THC gelatin-consuming animals, who ate significantly less gelatin than controls throughout the study, presumably due to the psychoactive effects of THC. A second benefit is that with this model, animals are able to remain in their home cage, minimizing stress of different environments and allowing normal consumption of chow and water. Third, the preparation of drug is straightforward, and does not require additional equipment for drug delivery aside from a gelatin cup. Additionally, it is possible to add multiple drugs to one gelatin, to examine drug combinations. Finally, oral consumption is a popular ingestion method for cannabis products, as it does not require vapor inhalation which could be aversive or ill-suited to certain populations (e.g. lung cancer patients), and drug effects last longer due to the pharmacokinetics of oral administration (Ohlsson et al, 1980). Our model is therefore a good preclinical representation of a major form of cannabis intake in humans. However, it is important to note that we did not observe escalation in this model with morphine gelatin, even when tolerance developed, which is something we would predict from a human taking morphine for pain (Zernig et al. 2007). Further testing with opioids like fentanyl or heroin may help to determie if escalation is possible with free-access gelatin.

In this study we developed a new device to measure real-time changes in gelatin mass. The piezoelectric load cell enabled us to record both the rapid changes in mass associated with animal contact, and cumulative changes in gelatin mass. By using this system, we were able to verify both the lower intake of THC that we first observed with our daily gelatin measurements, and an increase in consumption during the dark cycle compared to the light. Additionally, we observed gelatin consumption during every hour that was recorded, supporting a gradual intake of the gelatin suggested by our serum measurements. The total cost of components for the load cell measurement system is approximately $70, and with the schematics and source code freely available, it could be relatively easy to adapt this technology to other consumption media, including liquids (e.g. water, ethanol, etc.) or foods (e.g. chow, sucrose pellets).

The inclusion of ultrasonic clicks in our study was intended to capture an objective, ethological feature of the pain response; one that did not require an experimenter-determined score. This was partially driven by the fact that determining a pain response in animals (as in humans) is necessarily subjective, by way of a decision as to whether an animal made a specific motor movement (e.g. a paw withdrawal) or having to pick from several (e.g. hot plate jump, or paw lick, Williams and Porsolt, 2007). The observation that neuropathic pain produced increases in ultrasonic clicks, reversible by cannabinoids, suggests these clicks may represent either a painful or a general aversive state. Whether a click carries relevant con-specific information or is an epi-phenomenon of some other process, is unknown. One hypothesis is that a brief, broad spectrum click would be harder to audibly triangulate than a long call, and so mice could use clicks as a way of communicating a dangerous, aversive situation, without alerting nearby predators. Further ethological validation of this vocalization is warranted.

In conclusion, we have demonstrated that our gelatin self-administration paradigm allows voluntary consumption of drug over long periods, and may be useful as an alternative to classical operant behavioral designs for drug intake. Additionally, we have shown that mice produce click vocalizations that track pain status, which can be recorded non-invasively and may provide an objective correlate of pain. Finally, we have shown that the cannabinoids THC and CBD, in contrast to morphine, produce long-lasting relief of chronic neuropathic pain in a sciatic nerve injury model in mice. Specifically, CBD may represent a viable therapeutic option because of its low psychoactive profile, lack of efficacy in a pain-free state, and long-lasting reduction in allodynia in chronic pain.

## Acknowledgments

We thank the Mass Spectrometry facility at the University of Washington for serum processing.

Author contributions
EJYL, ADA, LCK, JJC, and BBL designed the study; BAW, ADA, and BBL designed the gelatin measurement apparatus; EJYL, ADA, and BAW performed research; EJYL, ADA, BAW, and BBL analyzed data; EJYL, ADA, and BBL wrote the manuscript.

