## Supplemental materials for "Orally consumed cannabinoids provide long-lasting relief of allodynia in a mouse model of chronic neuropathic pain"

* equal authorship

3 Supplemental Figure legends

3 Supplemental Figures

#### Supplemental Figure Legends

**Supplemental Figure 1.** Schematic of gelatin paradigm for drug gelatin pre- and post- sciatic nerve ligation. Blue-striped boxes indicate 2-day recording periods that are quantified and displayed in **Figure 1d**.

**Supplemental Figure 2**. **A**) Pictures of real-time recording device, annotated. **B**) Average number of feeding bouts per 2 hrs in the light vs dark cycle for each gelatin. Dark cycle consumption is significantly greater than light cycle for all groups (n = 3-4 per group). * $p < 0.05$

**Supplemental Figure 3**. **A**) Example of 2 non-clicks, which were rejected because of significant power in the audible range. **B**) The number of clicks in 5 minute recordings before and after pain testing for pain-naïve mice. Additionally, recordings from an empty cage show very few clicks in the absence of mice (n = 4 recordings for empty cage, n = 6 pain-naïve testing). * $p < 0.05$.

**Supplemental Figures**

### Supplemental Figure 1

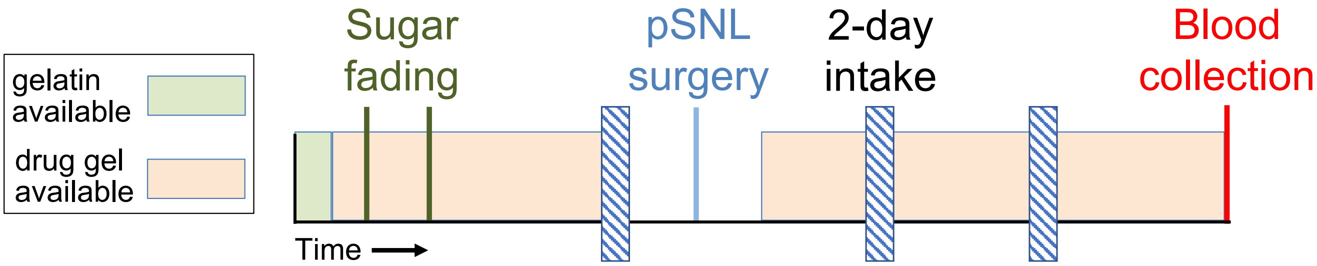

#### Supplemental Figure 2

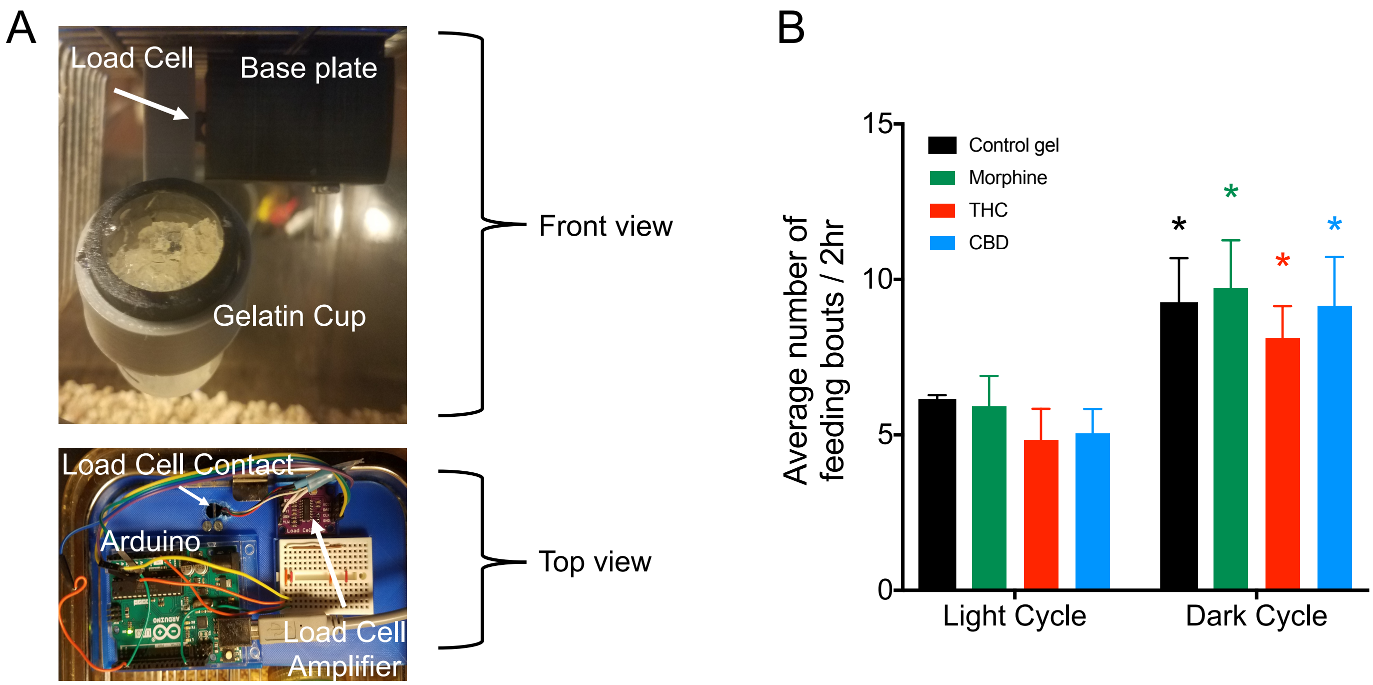

#### Supplemental Figure 3

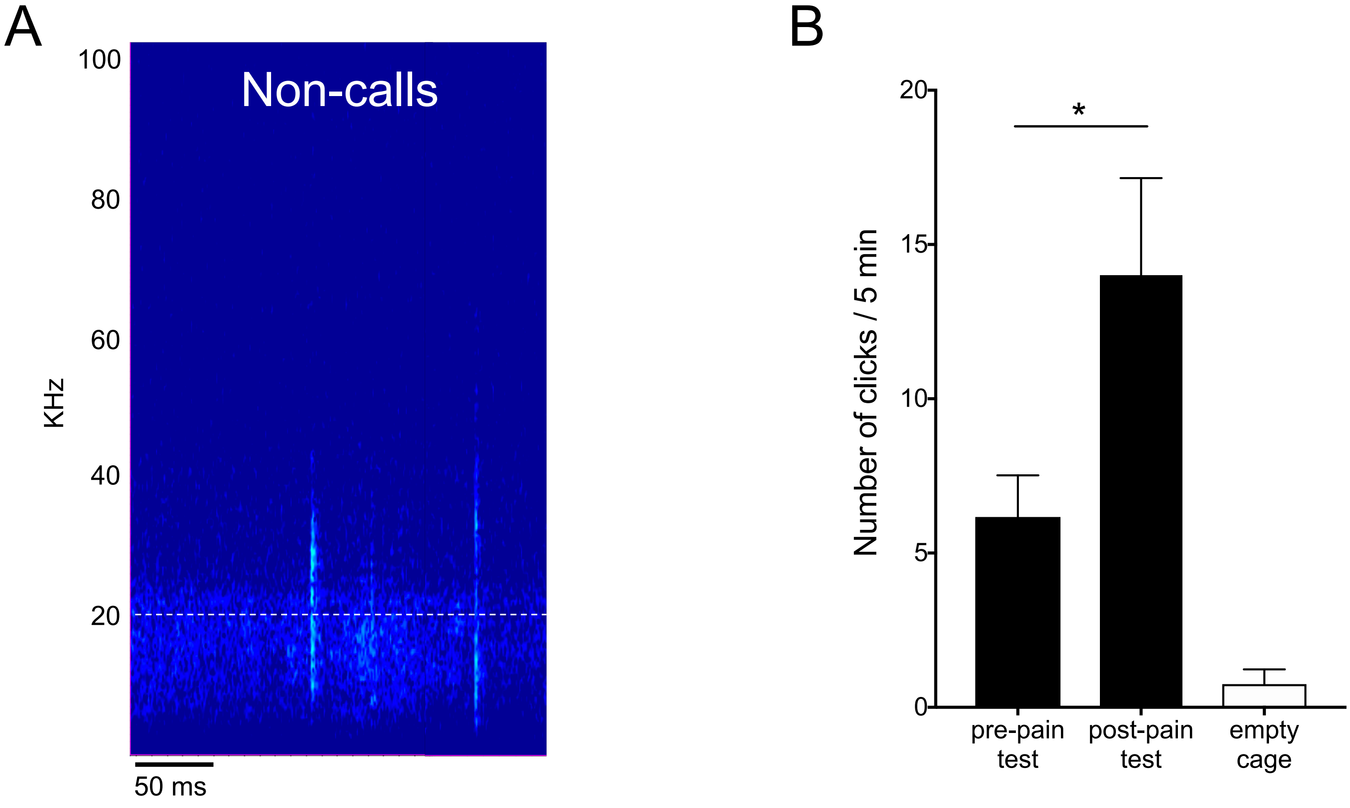
